# VaxLLM: Leveraging Fine-tuned Large Language Model for automated annotation of *Brucella* Vaccines

**DOI:** 10.1101/2024.11.25.625209

**Authors:** Xingxian Li, Yuping Zheng, Joy Hu, Jie Zheng, Zhigang Wang, Yongqun He

## Abstract

**Background:** Vaccines play a vital role in enhancing immune defense and preventing the hosts against a wide range of diseases. However, research relating to vaccine annotation remains a labor-intensive task due to the ever-increasing volume of scientific literature. This study explores the application of Large Language Models (LLMs) to automate the classification and annotation of scientific literature on vaccines as exemplified on *Brucella* vaccines.

**Results:** We developed an automatic pipeline to automatically perform the classification and annotation of *Brucella* vaccine-related articles, using abstract and title. The pipeline includes VaxLLM (Vaccine Large Language Model), which is a fine-tuned Llama 3 model. VaxLLM systematically classifies articles by identifying the presence of vaccine formulations and extracts the key information about vaccines, including vaccine antigen, vaccine formulation, vaccine platform, host species used as animal models, and experiments used to investigate the vaccine. The model demonstrated high performance in classification (Precision: 0.90, Recall: 1.0, F1-Score: 0.95) and annotation accuracy (97.9%), significantly outperforming a corresponding non-fine-tuned Llama 3 model. The outputs from VaxLLM are presented in a structured format to facilitate the integration into databases such as the VIOLIN vaccine knowledgebase. To further enhance the accuracy and depth of the *Brucella* vaccine data annotations, the pipeline also incorporates PubTator, enabling cross comparison with VaxLLM annotations and supporting downstream analyses like gene enrichment.

**Conclusion:** VaxLLM rapidly and accurately extracted detailed itemized vaccine information from publications, significantly outperforming traditional annotation methods in both speed and precision. VaxLLM also shows great potential in automating knowledge extraction in the domain of vaccine research.

**Availability:** All data is available at https://github.com/xingxianli/VaxLLM, and the model was also uploaded to HuggingFace (https://huggingface.co/Xingxian123/VaxLLM).

## Introduction

Vaccines utilizing the pathogens as whole or parts stimulate the immune system to generate memory-based immune cells to make a fast and robust immune response and kill the pathogens [1]. They have helped humans to reduce the infection of many diseases. Example successful vaccines include the HPV vaccine, MMR vaccine, COVID-19 vaccines, and many more [2]. The importance of vaccines against infectious disease has been clearly illustrated by the role of COVID-19 vaccines in saving millions of lives during the recent COVID-19 pandemic.

Among infectious diseases, zoonotic diseases pose unique challenges as they involve pathogens transmitted from animals to humans. For example, brucellosis is a zoonotic disease caused by *Brucella* spp., commonly transmitted to humans through contact with domestic animals such as cattle, sheep, and pigs [3]. *Brucella* vaccines, designed specifically to protect against *Brucella* spp., are primarily used in animals to prevent the spread of brucellosis within livestock and further reduce the risk of humans getting brucellosis [4,5]. Some commonly used *Brucella* vaccines include the *Brucella abortus* vaccine strain RB51, strain 19, and *Brucella melitensis* Rev.1 live attenuated vaccines [6]. Despite the risk to human health it causes, there is currently no approved human vaccine for brucellosis, underscoring the urgent need for research to accelerate vaccine development.

Some structured knowledgebase enhances our understanding of vaccines and provides comprehensive vaccine information. The VIOLIN vaccine knowledgebase [7,8] has so far included 4,708 vaccines for 217 pathogens or non-infectious diseases (e.g., cancer). The vaccines annotated in VIOLIN include licensed vaccines, vaccines being tested in clinical trials, and vaccines that have been studied in research and experimentally verified effective in at least one laboratory animal model. For each vaccine, VIOLIN has the annotated different types of vaccine knowledge, including vaccine name, platform, vaccination route, animal tested, etc. The knowledge in VIOLIN has also been represented using the Vaccine Ontology (VO), which is a community-based ontology model developed to include various aspects of vaccines, allowing clear hierarchical representation and specific vaccine querying, etc [9]. All the vaccines included in VO or VIOLIN are manually annotated from approximately 5,000 peer-reviewed articles. While 60 *Brucella* vaccines are included in the VIOLIN database, a plenty of experimentally verified *Brucella* vaccines have not been annotated and stored in VIOLIN. The manual collation method used in VIOLIN development requires experts reviewing to accurately extract detailed information about vaccine formulations, host species, and etc.. However, due to the rapid increase in the scientific literature and ongoing experiments with vaccines, this manual annotation process has become more challenging, as the volume of information quickly exceeds the ability of humans to review.

To address this challenge, the use of Large Language Models (LLMs), can significantly increase the speed of the acquisition of vaccine information [10]. Many LLM tools have been developed, such as PubTator [11] and Llama 3 [12] which both offer significant advantages in speeding up the speed and accuracy of scientific paper annotation. PubTator is widely used for biomedical text mining as it provides an API interface for annotations on key concepts such as proteins, genes, species, diseases, and chemicals based on Name Entity Recognition (NER). However, it does not provide the function of vaccine annotations.

On the other hand, Llama 3 (8 billion and 70 billion parameters) is an open source and free LLM developed by Meta. It has enhanced capabilities like a 128K token vocabulary and improved reasoning performance [13]. Llama 3 can be further fine-tuned to improve their ability to perform some specific tasks in the biomedical field. Instruction-supervised fine-tuning (Instruct Tuning) is one of the fine-tuning methods to optimize a model’s performance on generating responses that align precisely with user intentions. Instruction-supervised fine-tuning is done by having the model learn detailed instructions and the corresponding responses. To perform fine-tuning, LLaMA Factory [14] is a platform to streamline the fine tuning processes by offering a unified framework for fine-tuning LLM models, including the latest Llama 3.

In this manuscript, we report our development of a pipeline to systematically perform classification and standardized annotation of *Brucella* vaccine articles using a fine tuned VaxLLM (Vaccine Large Language Model). VaxLLM is derived from a Llama 3 8B model performed with an instruction-based fine tuning with a manually prepared dataset obtained from PubMed and VIOLIN. The whole pipeline contains systematic literature mining from PubMed, paper classification based on the amount of information extracted, and itemized paper annotation for filtered papers. PubTator was adopted to extract the host species information for cross-comparison and the standardized host gene information for gene enrichment analysis.

## Methods

### VaxLLM design and overall workflow

The overall VaxLLM design was shown in **Figure 1**. We first fine-tuned the Llama 3 model using the training data from VIOLIN and PubMed (purple part in Figure 1), which was then used to systematically process the articles (blue part). The paper processing starts with obtaining the “*Brucella* vaccine” articles by literature mining from PubMed. We then used the fine-tuned Llama 3 model to first classify the articles to filter the relevant articles containing specific information about *Brucella* vaccine. If the article is classified as “yes”, the fine-tuned Llama 3 model was then used to annotate these *Brucella* vaccine articles, specifically capturing key vaccine properties such as the platform, antigen, formulation, host species used as animal models, and the experimental methods employed to investigate the vaccine. The result was an Excel sheet compiling all the annotated information, providing a comprehensive dataset for further analysis. The Pubtator was also adopted to extract the biomedical entities such as genes, diseases, chemicals, and species for the article is classified as “yes” (green part). We extracted all the host genes identified by PubTator to perform a DAVID gene enrichment analysis (green part). The detailed methods of each step will be described in below sections.

**Figure 1:**
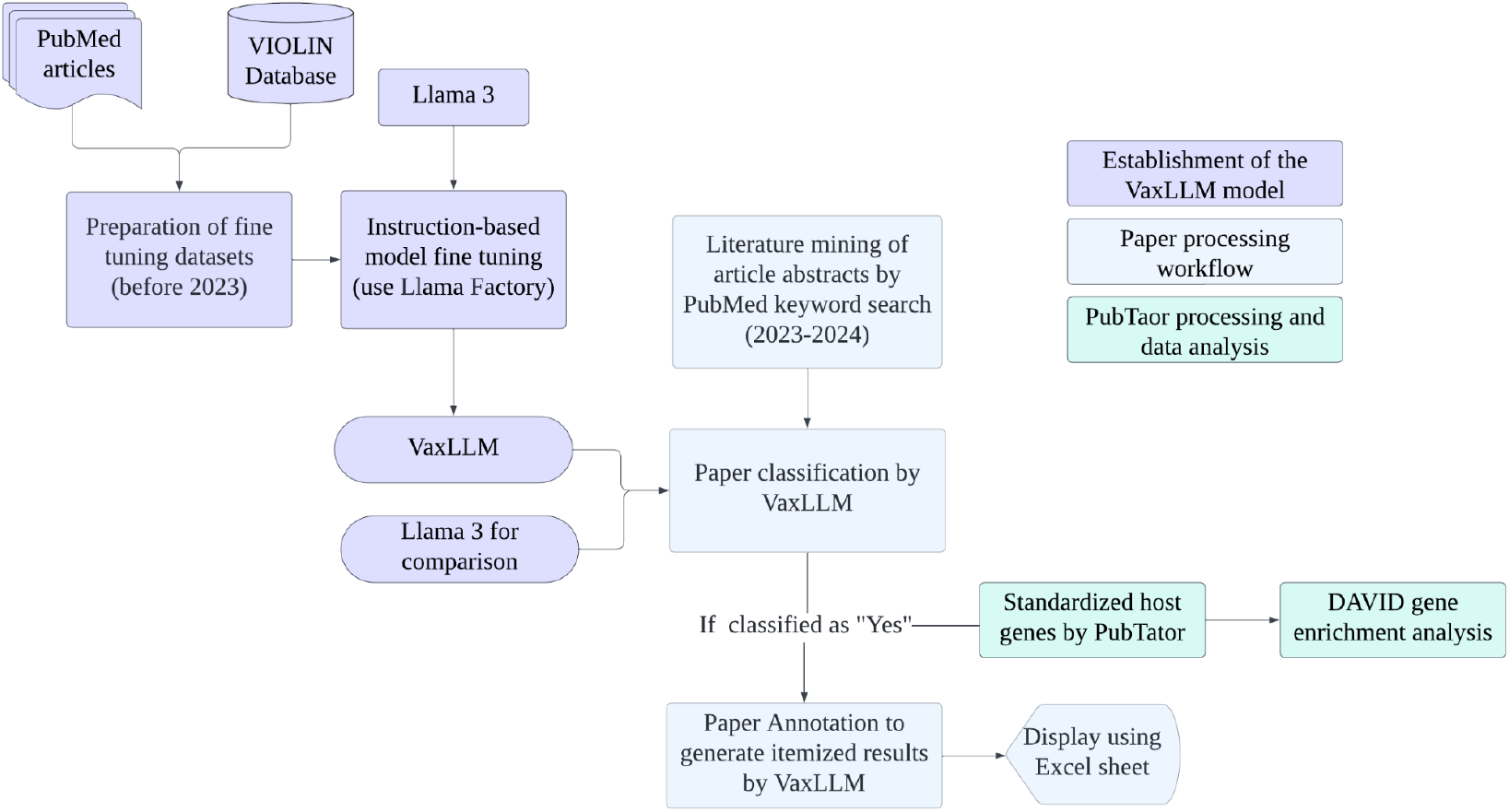
VaxLLM design and overall workflow.

### Llama 3 Model instruction based Fine-Tuning

We adopted the Llama 3 model with 8B parameters (https://ai.meta.com/blog/meta-llama-3/) for the classification and extraction of the *Brucella* vaccine information from PubMed abstracts. An instruction-based fine tuning was further done to the Llama 3 8B model to develop the VaxLLM pipeline for automated annotation of *Brucella* vaccine research articles.

The fine-tuning process consists of two parts: classification and annotation. The first was to train the model to classify a *Brucella* vaccine abstract and the second is to extract itemized information from the abstract. To create an effective classification training dataset, we compiled a collection of articles, consisting of 50 scientific article abstracts directly related to *Brucella* vaccines. This data is directly from the VIOLIN website (https://violinet.org/), which is used as the reference of the standardized annotation of 50 *Brucella* vaccines. We also prepared 100 negative examples from PubMed including 10 article abstracts not associated with *Brucella* vaccines, and 90 article abstracts related to *Brucella* but do not have specific *Brucella* vaccine information. Each article within the classification training dataset was labeled with a binary classification of “Yes” or “No” based on whether it can extract some information about specific *Brucella* vaccine formulation. The annotation training dataset includes 50 article abstracts from PubMed as the reference of the 50 *Brucella* vaccines from the VIOLIN website. We manually prepared the instruction based training dataset consisting of extracted itemized *Brucella* vaccine information, which includes ‘vaccine type’, ‘vaccine antigen’, ‘vaccine formulation’, ‘host species used as animal model’, and ‘experimental methods used to investigate vaccines’. The annotation information is based on the existing *Brucella* vaccine information provided by VIOLIN. The fine-tuning dataset preparation used the Alpaca format, which provides task-specific instructions to the model. Alpaca format includes the “instruction” as the prompt, “input” as the abstract of the articles, and “output” as the prepared correct responses for LLM to study.

The fine-tuning process adopted the Llama-Factory fine tune package, (https://github.com/hiyouga/LLaMA-Factory). The high memory efficient LoRA (Low-Rank Adaptation) fine tuning method and 4 bit quantization were used to reduce the memory requirement of hardware. Mixed-precision training (FP16) was also adopted to further reduce memory requirement and increase the speed. The learning rate was set to 5e-5 and the train batch size per device was set to 2 with gradient accumulation steps set to 4. After the fining tuning, the model is uploaded and available on HuggingFace named “VaxLLM” for later Usage.

### Systematically *Brucella* vaccine literature mining

We sourced relevant *Brucella* vaccine paper abstracts from PubMed, focusing on publications from 2023 to 2024 by conducting the “*Brucella* vaccine” term keyword search. It is noted that none of these papers published since 2023 was annotated in the existing VIOLIN system. To streamline the process, a Python script was adopted to perform the keyword search literature mining and extract essential data from each article, including the title, authors, DOI, and abstract. A total of 148 paper abstracts were identified from the previous process. The 148 paper abstracts were not overlapped with the training dataset.

### LLM prompt engineering and *Brucella* vaccine text extraction

We did prompt engineering for LLM to perform the specific task. This prompt helped us first identify whether the abstract provides available *Brucella* vaccine information or not. Then leverages the fine-tuned Llama 3 model to extract itemized information about the *Brucella* vaccine including “vaccine type”, “vaccine antigen”, “vaccine formulation”, “host species used as laboratory animal model”, and “Experiment Used to investigate the vaccine”. This prompt also gives a response template of each item, allowing better formatting for LLM.

The prompt used in our abstract information processing was divided into classification part and annotation part as follows:

~~~
   1. Vaccine Classification:
Using the following data: ‘**{Abstract information}**’, is this article about a
brucella vaccine? To classify an article as being about a brucella vaccine, you
must successfully extract at least some information about the vaccine formulation.
This includes details such as the antigen, protein, gene, adjuvant, or vaccine
platform mentioned in the abstract.
   2. Vaccine Annotations:
Extract the following details using the given data: ‘**{Abstract information}**’:
Vaccine Introduction,Vaccine Antigen, Vaccine Type, Vaccine Formulation, Host
Species Used as Laboratory Animal Model, Experiment Used to investigate the vaccine
Ensure each response is based solely on the provided data. Ensure the response is
formatted as follows:
Response:
Vaccine Introduction:
Vaccine Type:
Vaccine Antigen:
Vaccine Formulation:
Host Species Used as Laboratory Animal Model:
Experiment Used to investigate the vaccine:
~~~

The `{abstract information}` here is omitted. We used a Python script to automatically process the 148 abstracts in sections, where one abstract was processed at one time. We adjusted the max new token to be 200 to let LLM’s answer not be cut off in midstream. The ‘do sample’ is set to ‘FALSE’ in order for VaxLLM to respond to the provided abstract.

#### *Brucella* vaccine data representation

We developed a Python script to clean the VaxLLM output and transform it into an Excel spreadsheet for better visualization. This script processes each output entry, removes extraneous characters or errors, and ensures consistency across all annotation fields. The Excel spreadsheet contains the following columns for each of the 148 papers: PMID, Title, Authors, DOI, Citation, Abstract, Classification (“yes” or “no”), Vaccine Introduction, Vaccine Type, Vaccine Antigen, Vaccine Formulation, Host Species Used as Laboratory Animal Model, and Experiment Used to Investigate the Vaccine.

#### *Brucella* vaccine Name Entity Recognition using Pubtator

PubTator 3.0 has an API utilized to recognize the entity relations and synonyms to provide advanced search capabilities and enable large-scale analyses, streamlining many complex information needs. The Name Entity Recognition (NER) function of Pubtator also reads the paper abstract information. We used PubTator’s NER Recognition function to extract biomedical entities such as genes, proteins, mutations, species, and chemicals related to *Brucella*. We used a Python code to systematically provide the PMID to Pubtator and extract all the species information for each abstract for later cross comparison with VaxLLM extracted ‘host species used as laboratory animal model’. We also focused on the host genes extracted by Pubtator by systematically extracting all the host genes identified from PubTator and generating a host gene list.

### VaxLLM evaluation

We calculated the Precision, Recall, and F1-score to evaluate the performance of VaxLLM for classification tasks [15]. Precision means the proportion of correct predictions (true positive) out of all predictions and the higher precision indicates the higher accuracy of the model. Recall calculates the proportion of number of articles correctly classified as “yes” (true positive) out of number of articles correctly classified as “yes” (true positive) and number of articles incorrectly classified as “no” (false negatives). Recall reflects the model’s ability to identify relevant instances in a comprehensive way [16]. The F1 score provides a balanced measure that takes into account both precision and recall. The formula to calculate Precision, Recall, and F1-score is shown here:

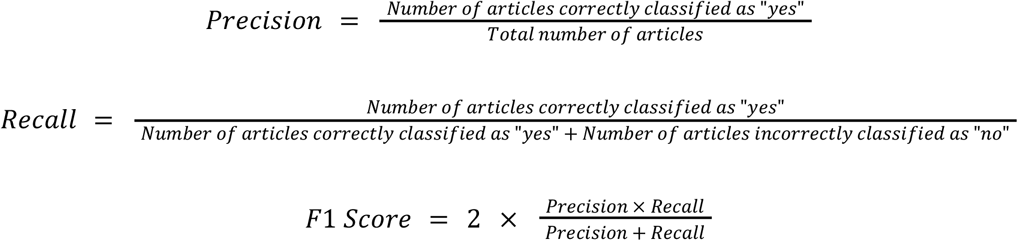

To evaluate the accuracy of VaxLLM’s annotations, we examined the following annotation categories: vaccine type, vaccine antigen, vaccine formulation, host species used as laboratory animal model, and experiment used to investigate the vaccine. We adopted human manual checking as the gold standard and calculated the accuracy to be the proportion of correct annotations out of the total number of annotated articles.

### Data access and display

*Brucella* vaccine data and documentation is available at GitHub: All source codes are available at GitHub: https://github.com/xingxianli/VaxLLM. The model was also uploaded to HuggingFace (https://huggingface.co/Xingxian123/VaxLLM).

## Results

### *Brucella* vaccine data collection

For the 148 articles retrieved from PubMed, we manually classified and annotated these articles. Those *Brucella* vaccine data were used as the testing data for VaxLLM. To classify an article as “Yes” in the gold standard dataset, the article is required to include specific vaccine formulation. Among the 148 articles, 58 papers were classified as “Yes” and 90 classified as “No”. We also manually annotated and extracted the “vaccine antigen”, “vaccine platform”, “vaccine formulation”, “Host species used as animal models”, and “experiments used to investigate the vaccine” based on the abstract information of each article. “Vaccine Platform” and “Host species used as animal models” were used as examples of the coverage of the gold standard dataset (**Table 1**), which demonstrates the inclusion of various vaccine platforms and tested host species in the dataset.

**Table 1.** Statistics of annotated *Brucella* vaccines as gold standard for VaxLLM development.

|  | Overall |
| --- | --- |
| **Vaccine Platform** | N=58 |
| Live attenuated vaccine | 26 (44.8%) |
| Subunit vaccine | 18 (31.0%) |
| RNA vaccine | 2 (3.4%) |
| Recombinant vector vaccine | 4 (6.8%) |
| Conjugate vaccine | 2 (3.4%) |
| Other / Not mentioned | 6 (10.3%) |
| **Host species used as animal model** | N=58 |
| Mice (including BALB/c and C57BL/6) | 28 (48.3%) |
| Cow | 8 (13.8%) |
| Sheep | 4 (6.9%) |
| Other / Not mentioned | 18 (31.0%) |

### Fine-Tuning: From Baseline Llama 3 to VaxLLM

We first adopted the baseline Llama 3 8B model (non-fine-tuned) on the same dataset to evaluate its performance in classifying and annotating the *Brucella* vaccine related information. However, the non-fine-tuned model produced incomplete and inconsistent results, particularly mentioning the “Not specify” in the annotation part even if the information is included. The non-fine tuned model also did not perform well on *Brucella* paper classification, by misclassifying “yes” articles to “no”, resulting in the great loss of vaccine information.

Recognizing these limitations, we fine-tuned the baseline Llama 3 8B model and developed the VaxLLM model. VaxLLM specialized in accurate classification and detailed annotation of the *Brucella* vaccine related information. The model was uploaded to the HuggingFace (see Methods), allowing direct usage by the public.

### VaxLLM Classification Performance Analysis

To evaluate the performance of the VaxLLM in classifying *Brucella* vaccine articles, we used human classification as the gold standard for accuracy. Out of 148 papers in the gold standard dataset, VaxLLM correctly classified 60 articles as “yes” (true positives) and 84 as “no” (true negatives), while misclassifying 7 articles as “yes” that should have been “no” (false positives). Notably, VaxLLM produced no false negatives. We used the accuracy, precision, recall, and F1 score to calculate the performance of the VaxLLM classification task. VaxLLM has an accuracy of 0.95, precision of 0.90, recall of 1.0, and F1_score to be 0.95.

To better compare the difference before and after the fine tuning process, we used the confusion matrices (**Figure 2)** to summarize the classification results for both the fine-tuned VaxLLM model and the non-fine-tuned baseline Llama3 8B model. Each matrix shows four categories: True Negatives (top-left), False Positives (top-right), False Negatives (bottom-left), and True Positives (bottom-right). The fine-tuned model (left) achieved 84 True Negatives, 7 False Positives, 0 False Negatives, and 60 True Positives, reflecting a higher classification precision and recall. In contrast, the non-fine-tuned model (right) resulted in 60 True Negatives, 29 False Positives, 3 False Negatives, and 56 True Positives, indicating comparatively lower accuracy due to increased misclassifications.

**Figure 2.**
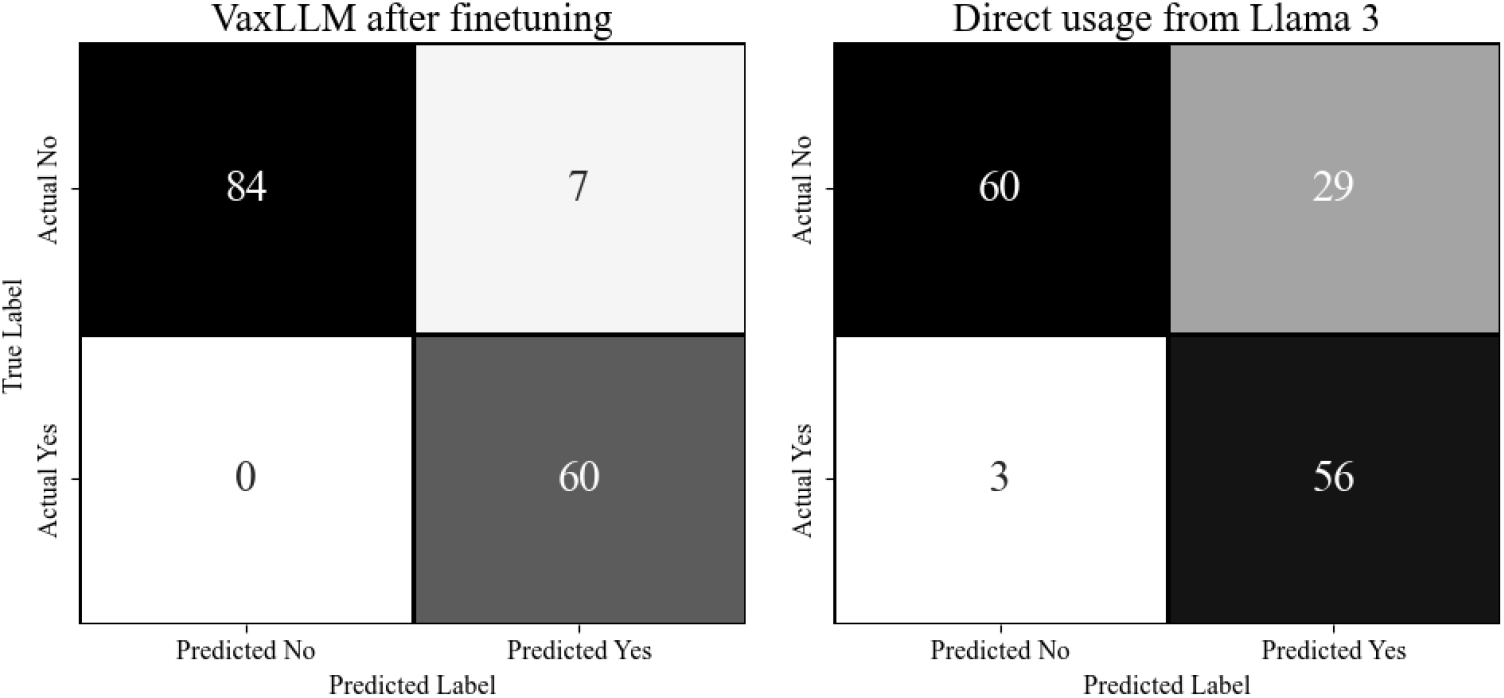
Confusion Matrix comparison of the VaxLLM and non-fine-tuned Llama 3 model.

A receiver operating characteristic (ROC) curve was drawn for the demonstration of fine-tuned Llama 3 ability of classifying the articles (**Figure 3)**. Specifically, the fine-tuned VaxLLM achieved the area under the ROC curve (AUC) of 0.71. Compared to the baseline Llama 3 model, which AUC is 0.63, VaxLLM consistently outperforms the non fine tuned model across a range of classification thresholds, demonstrating its ability to classify *Brucella* vaccine relevant articles.

**Figure 3.**
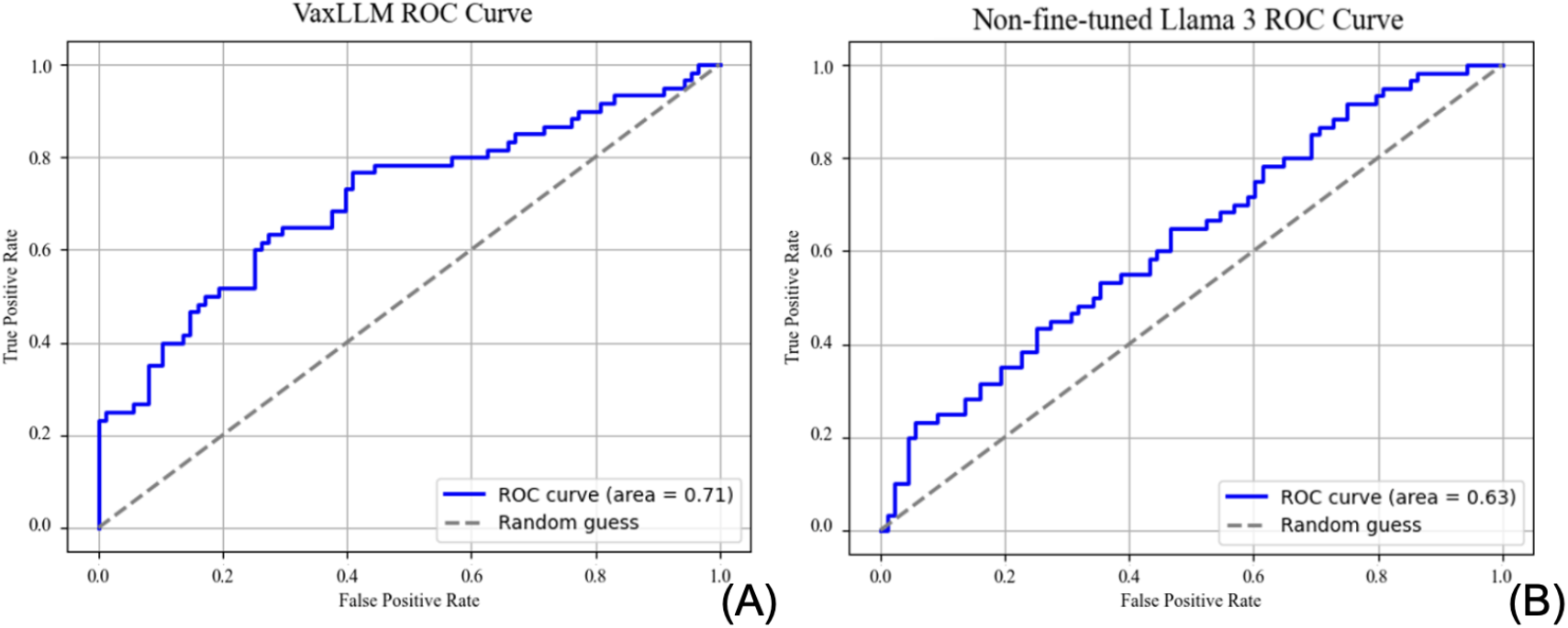
ROC curve of fine tuned VaxLLM (A) and the baseline Llama 3 model (B)

To further explore the reasons for the 7 false positive results, we surveyed these papers in more detail **(Table 2)** [17–23]. Specifically, these seven false positive papers include one review article, 2 articles focused on mechanism understanding, and 4 articles focussed on *Brucella* diagnosis instead of vaccine development. These were indeed not focused on *Brucella* vaccine development.

**Table 2.** Details of the 7 false positive articles by VaxLLM.

| PMID | Article title | Overview | Possible misclassifying reason |
| --- | --- | --- | --- |
| 39151885 | The development of a human *Brucella* mucosal vaccine: What should be considered? | Reviews immune mechanisms and strategies for mucosal vaccine development without describing a specific vaccine formulation. | Review paper of *Brucella* mucosal vaccines |
| 39024932 | Establishment of an A/T-Rich Specifically MGB Probe digital droplet PCR Assays Based on SNP for *Brucella* wild strains and vaccine strains. | Discusses development of diagnostic tools for identifying *Brucella* strains rather than a study of the vaccine itself. | Diagnostic assay focused; contains some *Brucella* protein |
| 38352039 | Fecal and vaginal microbiota of vaccinated and unvaccinated pregnant elk challenged with *Brucella* abortus | Explores microbiota changes in vaccinated elk but does not investigate or detail the vaccine formulation. | Indirect vaccine effects |
| 38101579 | Use of recombinant malate dehydrogenase (MDH) and superoxide dismutase (SOD) [CuZn] as antigens in indirect ELISA for diagnosis of bovine brucellosis. | Focuses on diagnostic testing using antigens rather than vaccine formulation or effects. | Focus on diagnostic tools, contains some *Brucella* antigens |
| 38003334 | Regulation of the Gene for Alanine Racemase Modulates Amino Acid Metabolism with Consequent Alterations in Cell Wall Properties and Adhesive Capability in *Brucella* spp | Discusses mutant strains as potential vaccine candidates without a developed or tested vaccine formulation. | Mentions potential vaccine candidate but not developed |
| 37812358 | Serological, cultural, and molecular analysis of *Brucella* from Buffalo milk in various regions of Iran. | Mentions detection of the RB51 vaccine strain but focuses primarily on diagnostic methodologies. | Vaccine strain referenced indirectly |
| 37389214 | The outer membrane proteins based seroprevalence strategy for *Brucella* ovis natural infection in sheep | Discusses detection of specific antigens (OMP25 and OMP31) for diagnostics rather than their role in a vaccine. | Mentions antigens for antigen detection and *Brucella* diagnosis |

### VaxLLM output demostration

The result of identified *Brucella* vaccine itemized information using VaxLLM was exported to an Excel sheet. To further illustrate the effectiveness of VaxLLM annotation, we present a cleaned output example in **Figure 4** (Panel B), demonstrating the results for paper PMID 39220468 [24].

**Figure 4.**
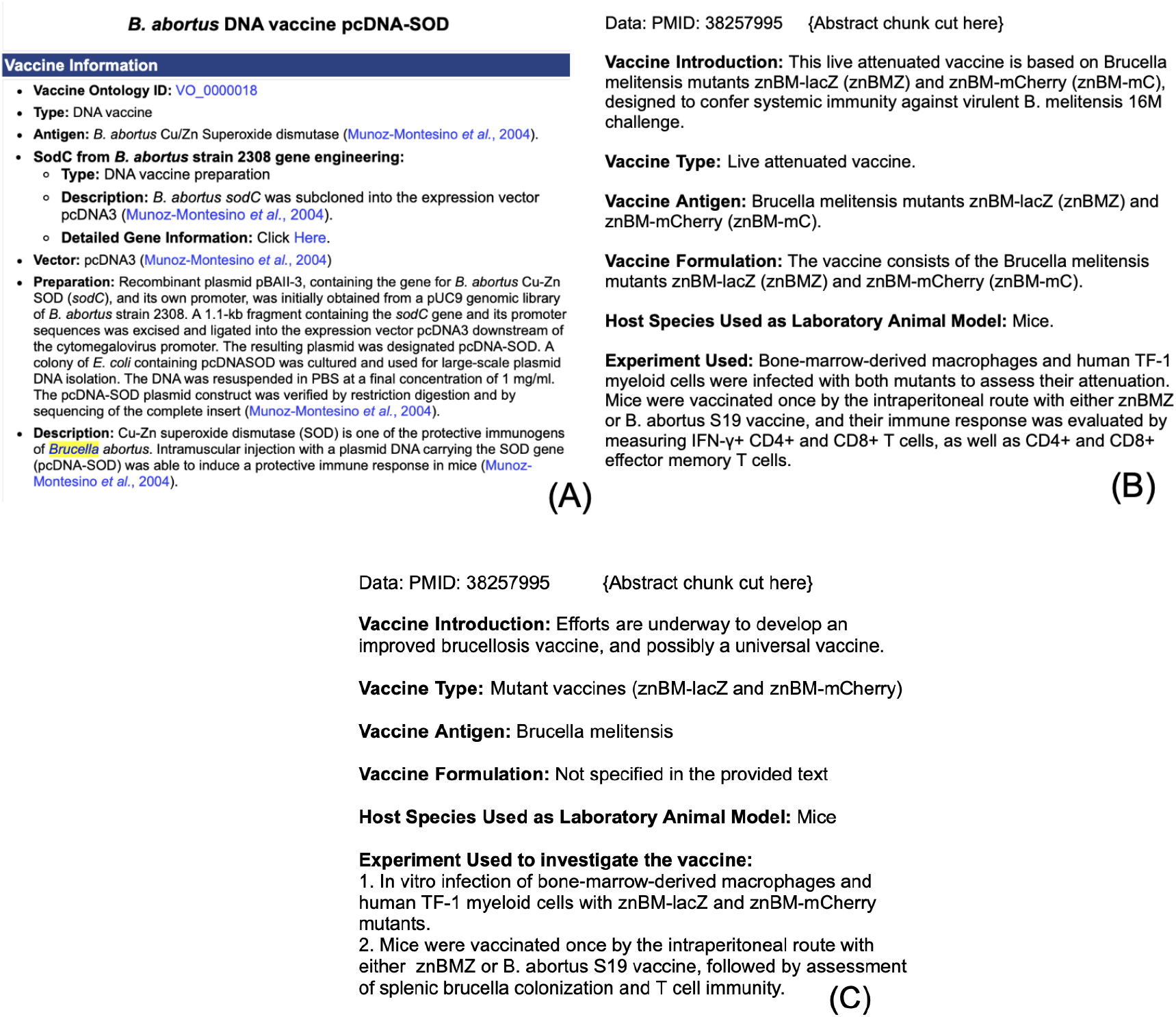
VaxLLM annotation output example compared to VIOLIN and Llama 3 8B. (A) VIOLIN report screenshot. (B) VaxLLM after fine tuning output example usage for paper PMID 38257995. (C) Llama 3 8B direct usage for paper PMID 38257995

**Figure 4** compares the VaxLLM output (Panel B) with a VIOLIN report screenshot (Panel A), highlighting VaxLLM’s ability to produce similarly structured vaccine annotations. The (Panel C) includes the same article annotated by baseline Llama 3 8B model, which includes many incorrect annotations. VIOLIN was implemented as standardized annotations, with some already mapped into Vaccine Ontology (VO). Since VaxLLM was trained on data from VIOLIN, the outputs from VaxLLM are likewise relatively standardized. This similarity ensures that VaxLLM’s extracted information can be directly integrated into VIOLIN after a simple verification step, significantly reducing the time and effort compared to manual curation. The VaxLLM automation has the great potential to accelerate the integration of *Brucella* vaccine data in the VIOLIN database and enhance the efficiency of curating new research findings.

We also compared the accuracy of VaxLLM and the non-fine-tuned model in five key categories: Vaccine Type, Vaccine Antigen, Vaccine Formulation, Host Species Used as Laboratory Animal Model, and Experiment Used to Investigate the Vaccine (**Figure 5**). The overall accuracy of the VaxLLM model was significantly higher at 97.9% compared to 80.0% for the non-fine-tuned model.

**Figure 5.**
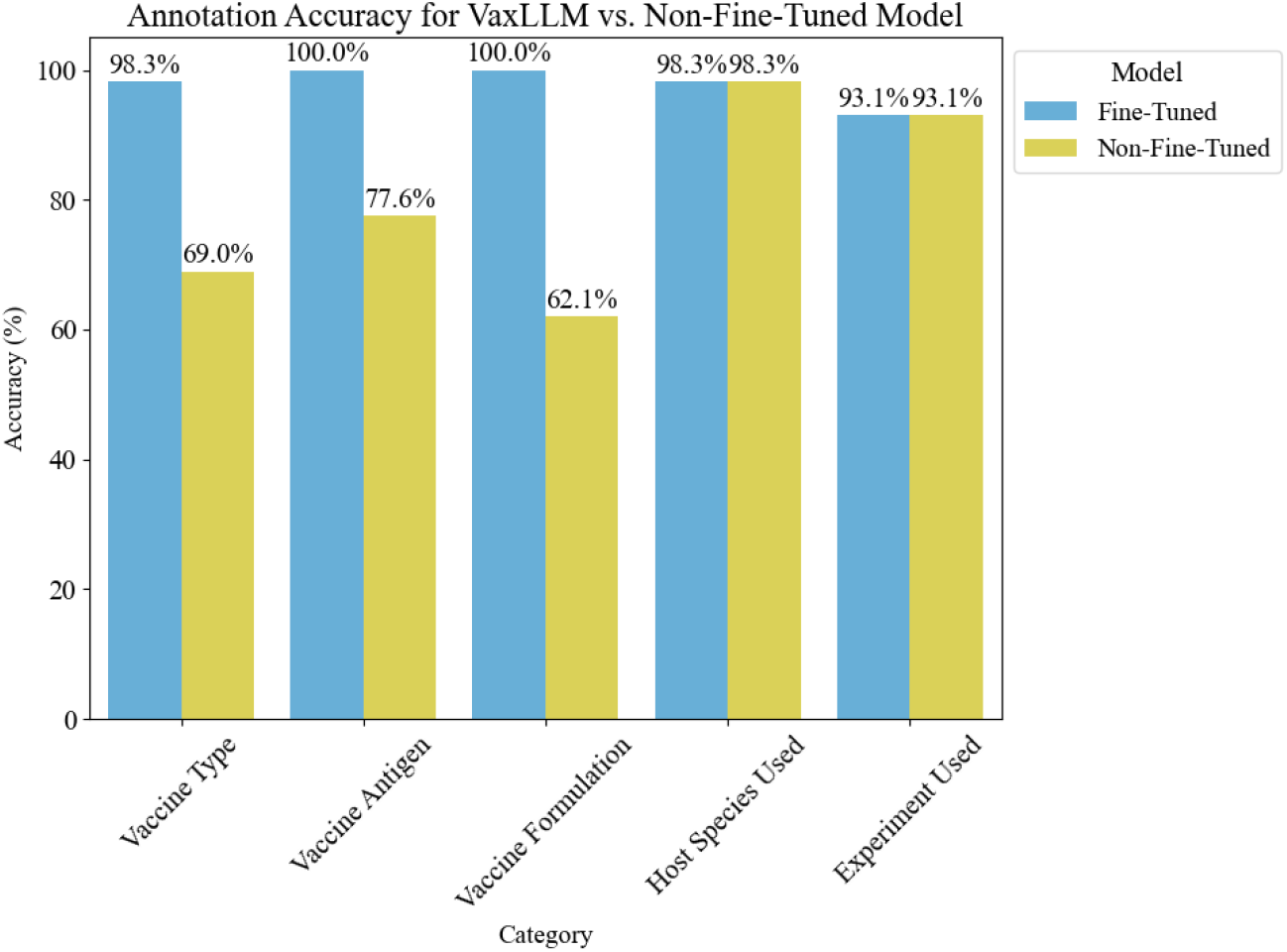
Bar chart comparing the annotation accuracy of VaxLLM and non-fine-tuned Llama.

In addition to the accuracy of VaxLLM, we found that the VaxLLM provided more standardized annotation output than non fine-tuned models. For example, the Vaccine type extracted by non-fine-tuned Llama 3 includes extraneous information making it difficult to standardize the result (ie. map to a standardized database) later on. For PMID 39218373 [25], VaxLLM identified the vaccine type as ‘subunit vaccine’, while the non-fine-tuning model identified the vaccine type to be ‘Subunit vaccine (contains multiple epitopes and IgV_CTLA-4)’. Moreover, for some standardized vaccine type terms such as ‘Live attenuated vaccine’, while VaxLLM identified it to be the standardized form, the non-fine tuned model will identify it as various names, including ‘Live-attenuated’, ‘Live bacteria-based vaccine’, ‘Live modified vaccine’, etc. The fine-tuning process entitles VaxLLM the ability to extract in a disciplined way because the fine-tuning dataset was prepared using VIOLIN database annotation, which is already in a formulated way. The standardized annotation of VaxLLM offers the convenience of standardizing the annotation in the future.

### PubTator usage for cross comparison and gene enrichment analysis

In our analysis of *Brucella* vaccine data, we integrated multiple tools to enhance the accuracy and depth of our annotations. VaxLLM classified 58 articles as relevant (“Yes”), for which we conducted detailed information extraction. This classification step ensured that subsequent analyses (e.g. PubTator-based entity extraction) were performed on a highly concentrated dataset, underscoring the significance of accurate classification in streamlining the annotation process. Using PubTator, we extracted standardized biomedical entities, such as genes, species, diseases, and chemicals, from these articles. From a total of 58 articles, we identified 54 unique host genes, which were then narrowed down to 37 genes standardized for DAVID gene enrichment analysis. We focused on terms with a False Discovery Rate (FDR) below 0.05 to identify statistically significant terms. **Figure 6** shows the bar chart of the top ten terms with the highest fold enrichment, annotated with the associated gene count for each term.

**Figure 6.**
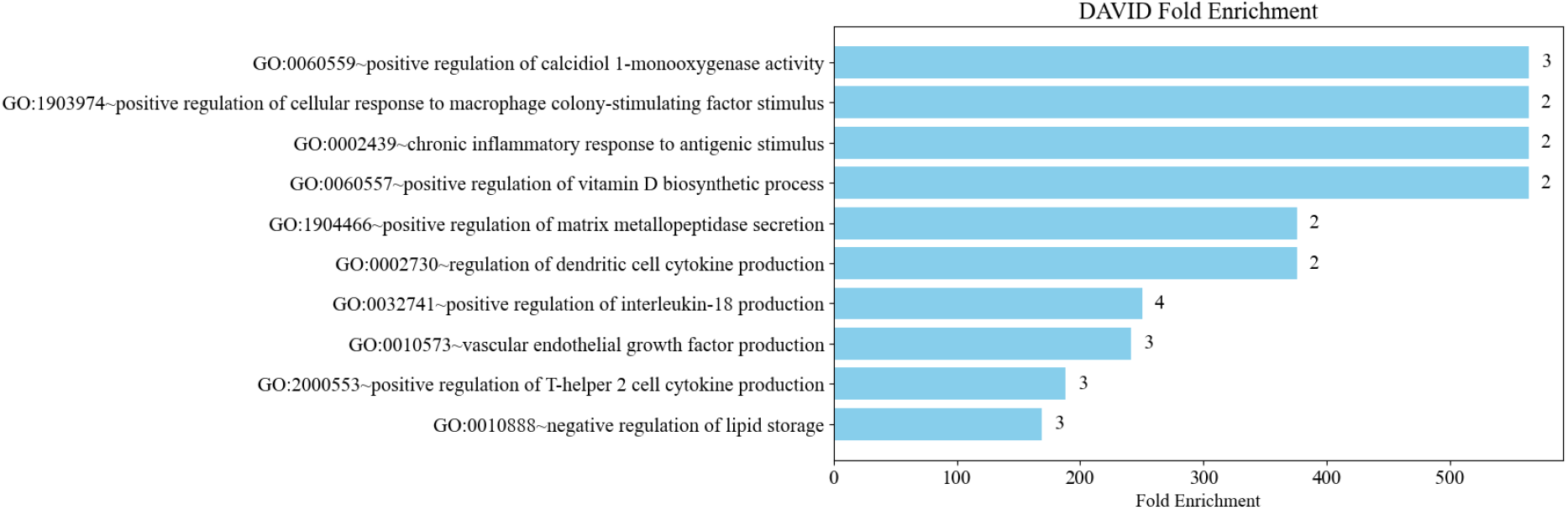
Gene Ontology (GO) gene enrichment results based on DAVID analysis.

Additionally, PubTator is a useful tool in identifying the biomedical entities in abstracts. We compared the PubTator extracted species information with “Host species used as laboratory animal models” identified by VaxLLM. VaxLLM and PubTator overlapped 45 instances as lab animal models, while PubTator missed 7 of these lab models. Compared to the human annotation as gold standard, only 3 of VaxLLM host species annotations were incorrect. By combining VaxLLM’s domain-specific classification capabilities with PubTator’s comprehensive biomedical entity extraction, we can maximize the precision of context-based annotations and VaxLLM’s ability for context based annotations.

## Discussion

Our pipeline aimed for automated vaccine information extraction from articles related to *Brucella* vaccines. By leveraging the fine-tuned capabilities of VaxLLM, we can automate the process from literature mining to extracting structured, itemized vaccine information. The structured and accurate output of VaxLLM lays a solid foundation when integrating those annotations into databases like VIOLIN. The annotated information can be rapidly reviewed and incorporated, significantly reducing the time and and effort required for manual collation in database development. The classification step in this pipeline introduces additional use cases, enabling researchers to conduct highly concentrated studies on *Brucella* vaccines.

Some existing approaches in the field of vaccine information extraction rely heavily on keyword-based methods [26,27]. While these methods show utility in information retrieval and abstract classification, they often lack the contextual understanding required to accurately label vaccine-related information. In contrast to these methods, VaxLLM provides structured vaccine-specific category information like platform, antigen, vaccine formulation, etc. based on the provided context. Particularly in the underexplored domain of Brucella vaccine research, VaxLLM’s capability to move beyond static keyword lists delivers flexible annotations tailored to specific vaccine research needs.

In future directions, a key aspect we consider is the development of an ontology-based knowledge graph for use in a Knowledge Graph Retrieval Augmented Generation (KG-RAG) [28] approach to graphs to improve VaxLLM’s ability. The KG-RAG can use the knowledge graph to structure vaccine specific data into nodes and relationships, enabling more effective query handling and information retrieval [29]. By converting VO into a graph-based structure, we can encode hierarchical and relational vaccine data, such as platforms, formulations, and host species, as interconnected nodes and edges [30]. Integration of this knowledge graph into VaxLLM through the KG-RAG enables standardized querying and retrieval. For instance, when the VaxLLM provides annotation information of “vaccine platform” for a specific *Brucella* strain, the KG-RAG approach could dynamically query the graph to retrieve standardized data, such as IDs in VO, along with related annotations.

*Brucella* vaccines were used as an example for the VaxLLM development and testing. We demonstrated the feasibility of VaxLLM and evaluated the performance of VaxLLM in the focused setting of *Brucella* vaccines. We also aim to expand the scope to include all vaccines in the VIOLIN database, which includes both infectious and non-infectious diseases (e.g. cancer vaccines). By including more vaccine types, VaxLLM would be generalized to a wide range of vaccines. Our pipeline could then assist in identifying emerging vaccine trends, analyzing research gaps, and providing actionable insights to researchers.

While VaxLLM provides relatively standard structured annotations, the limitation is the lack of integration with standardized databases. Therefore, another future direction is to integrate these results into bio-ontologies (e.g. Vaccine Ontology) and provide standardized IDs. Mapping the annotations into an ontology will allow for precise hierarchical querying of vaccine data and facilitate data integration between different studies and domains [31]. Such standardization will improve the consistency of the annotations and further improve the aid to research validity.

## Conclusions

Overall, our study demonstrates the potential of using large language models to rapidly and accurately annotate *Brucella* vaccine-related literature. We present an automated pipeline, powered by the VaxLLM model, to extract itemized vaccine information from scientific papers. The VaxLLM model was trained and subsequently evaluated for its ability to classify articles and provide detailed itemized annotations. This work accelerates the processing of new data, contributing to large-scale vaccine databases such as VIOLIN with high-quality, comprehensive annotations.

## Competing Interests

The authors declare that they have no competing interests.

## Funding

This research has been supported by the NIH U24 grant (U24AI171008).

## Author contributions

XL: VaxLLM development, *Brucella* vaccine data annotation, method evaluation, first version of manuscript writing; YZ: cleanup script development; JH: *Brucella* vaccine data annotation; JZ: data and result interpretation and discussion; ZW: LLM instruction, data and result interpretation and discussion; YH: Project design, *Brucella* vaccine domain expert, and data and result interpretation. All authors approved the paper publication.

## Acknowledgements

This research was supported by the National Institutes of Health grant U24AI171008.

## Notes

### Competing Interest Statement

The authors have declared no competing interest.

